# A rapid, super-selective method for detection of single nucleotide variants in *C. elegans*

**DOI:** 10.1101/2020.04.01.020818

**Authors:** Denis Touroutine, Jessica E. Tanis

## Abstract

With the widespread use of single nucleotide variants generated through mutagenesis screens, the million mutation project, and genome editing technologies, there is pressing need for an efficient and low-cost strategy to genotype single nucleotide substitutions. We have developed a rapid and inexpensive method for detection of point mutants through optimization of SuperSelective (SS) primers for end point PCR in *Caenorhabditis elegans*. Each SS primer consists of a 5’ “anchor” that hybridizes to the template, followed by a non-complementary “bridge,” and a “foot” corresponding to the target allele. The foot sequence is short, such that a single mismatch at the terminal 3’ nucleotide destabilizes primer binding and prevents extension, enabling discrimination of different alleles. We explored how length, stability, and sequence composition of each SS primer segment affected selectivity and efficiency in order to develop simple rules for primer design that allow for distinction between any mismatches in various genetic contexts over a broad range of annealing temperatures. Manipulating bridge length affects amplification efficiency, while modifying the foot sequence can increase discriminatory power. Flexibility in the positioning of the anchor enables SS primers to be used for genotyping in regions with sequences that are challenging for standard primer design. In summary, we have demonstrated flexibility in design of SS primers and their utility for genotyping in *C. elegans*. Since SS primers reliably detect single nucleotide variants, we propose that this method could have broad application for SNP mapping, screening of CRISPR mutants, and colony PCR to identify successful site-directed mutagenesis constructs.

## INTRODUCTION

In this genomic era, researchers have identified a multitude of single base pair substitutions, the most common type of DNA sequence variation in genome sequence data. Naturally occurring single nucleotide polymorphisms have been linked to human disease (Shastry 2002; Suh and Vijg 2005) and are used for gene mapping (Davis *et al*. 2005; Altshuler *et al*. 2008) and evolutionary studies (Koch *et al*. 2000). In genetic model systems, point mutants isolated through mutagenesis screens and gene editing are essential tools for discovery of gene function. Therefore, researchers working across a wide range of disciplines and systems can greatly benefit from having a low cost, robust, and efficient method to distinguish between alleles with single nucleotide variations.

In *C. elegans*, many mutants have been generated in forward genetic screens, with the most commonly used chemical mutagen ethyl methanesulfonate (EMS) exhibiting a mutagenesis bias towards transition mutations (Brenner 1974; Flibotte *et al*. 2010). Over 800,000 single nucleotide substitutions (SNSs) have been identified in the million mutation project, carried out to provide the *C. elegans* research community with a resource of mutant alleles for all genes in the genome (Thompson *et al*. 2013). SNSs are now also induced by CRISPR gene editing to interrogate the function of specific amino acids (Dickinson and Goldstein 2016). To analyze the phenotype associated with a mutation and decipher gene function, genetic crosses are performed, necessitating a reliable, rapid method for routine genotyping of SNSs.

A variety of techniques for SNS genotyping are available, however, these methods are either labor intensive, expensive, or require extensive troubleshooting (Mamotte 2006). Cleaved Amplified Polymorphic Sequence (CAPS) genotyping is based on the formation or disruption of a restriction enzyme recognition site by a mutation and involves enzymatic digestion of DNA amplified from the target region followed by electrophoresis (Konieczny and Ausubel 1993). A modified method, dCAPS, can be used to create or remove a restriction enzyme site to distinguish between two alleles (Neff *et al*. 2002). While the CAPS method is simple, it involves extra steps beyond PCR, requires purchase of the different restriction enzymes, and can lead to ambiguous results in cases of incomplete enzyme digestion. Other genotyping methods, including the TaqMan assay and melting curve analysis of FRET probes, are not labor intensive, but do require acquisition of allele-specific hybridization probes labeled with different fluorescent dyes as well as access to expensive instrumentation to allow for real time monitoring of PCR amplification (Bernard *et al*. 1998; Livak 1999).

Allele-specific PCR, also known as Amplified Refractory Mutation System (ARMS) PCR, and the modified method Simple Allele-discriminating PCR (SAP) are inexpensive genotyping methods which utilize allele-specific oligonucleotide primers (Newton *et al*. 1989; Little 2001; Bui and Liu 2009; Medrano and De Oliveira 2014). Discrimination between wild-type and mutant alleles is based on a mismatch at the 3’ terminal base which prevents extension of the primer (Petruska *et al*. 1988; Newton *et al*. 1989; Wu *et al*. 1989; Huang *et al*. 1992). However, ARMS and SAP often require extensive troubleshooting as PCR specificity must be controlled by stringent reaction conditions. Further, a lack of flexibility in primer placement can make SNS detection difficult in some genetic contexts (Medrano and De Oliveira 2014).

To detect the presence of rare single nucleotide polymorphisms in DNA fragments found in blood samples, Vargas et al. (2016) developed SuperSelective (SS) primers for real-time PCR assays. A SS primer consists of a 5’ anchor sequence that hybridizes to the template DNA followed by a non-complementary bridge sequence and a short 3’ foot sequence that is complementary to the target allele sequence (Vargas *et al*. 2016). Our goal was to design and optimize allele-specific primers for end point PCR genotyping based on the principle of SS primers. We have probed the different regions of the primer to determine how specificity is achieved and developed simple rules for SS primer design. Our work presents SuperSelective genotyping as an advantageous alternative to existing genotyping methods that will facilitate research with genetic systems.

## MATERIALS AND METHODS

### Nematode Culture

*C. elegans* were maintained on Nematode Growth Media (NGM) plates with OP50 *E. coli* as a food source using standard techniques. The wild-type strain was Bristol N2. Strains and alleles used in this study were as follows: PT443 *klp-6(sy511)* III; *him-5(e1490)* V; DM1017 *plx-2(gk2864)* II, *C05B5*.*11(gk2895)* III; VC40549 *cil-7(gk688330)* I; ZZ12 *lev-11(x12)* I; CB1372 *daf-7(e1372)* III; DA465 *eat-2(ad465)* II. All strains were maintained at 20°C except CB1372 which was grown at 15°C.

### Molecular biology

Primers were designed as described in the results section and obtained from Integrated DNA Technologies (IDT). A complete list of primers used is in Supplementary Table 1. Genomic DNA (gDNA) was isolated with the Gentra Puregene Tissue Kit (Qiagen Cat. No. 158667) following the manufacturer’s instructions. Crude genomic DNA was extracted by incubating worms in lysis buffer (50 mM KCl, 10mM Tris pH 8.3, 2.5 mM MgCl2, 0.45% NP-40, 0.45% Tween-20 and 1 mg/ml of Proteinase K) for 1 hour at 65°C followed by 95°C for 25 minutes and used where indicated. PCR was performed in 15 µl reactions using 2x GoTaq DNA Polymerase master mix (Promega Cat. No. M3008) with 5 ng of gDNA and 500 nM of each primer. The following protocol was performed: 98°C for 30 seconds (cycle one only), 98°C for 10 seconds, annealing temperature (gradient) for 15 seconds, and 72°C for 30 seconds for 30 cycles. Annealing temperatures for the anchor of SS primers were determined using the New England Biolabs T_m_ calculator and are indicated in Supplementary Table 1. Gradient temperatures were across 10°C; T_m_ minus 5°C (lowest temperature) to T_m_ plus 5°C (highest temperature). PCR products were resolved on 1% agarose gels and visualized with SYBR safe (Thermo Fisher Cat. No. S33101). All gradient PCR experiments were performed at least twice; representative images displayed in the figures.

### Data availability

All data and methods required to confirm the conclusions of this work are within the article, figures, and table.

## RESULTS

### Limitations of ARMS PCR for *C. elegans* genotyping

We have been performing genetic crosses with point mutants that do not cause visible phenotypes or change restriction sites for multiple ongoing projects. To discriminate between wild-type and mutant alleles we sought to use the ARMS PCR genotyping strategy, which is based on the principle that a mismatch at the 3’ terminal base of a primer results in inefficient amplification (Petruska *et al*. 1988; Wu *et al*. 1989; Huang *et al*. 1992) as the absence of exonuclease activity in Taq DNA polymerase prevents primer-template mismatch repair (Tindall and Kunkel 1988). We designed two allele-specific forward primers that hybridized to the variant base in either the wild-type or mutant template. Each allele-specific primer was paired with a common reverse primer and gradient PCR reactions were performed to determine the optimal temperature for discriminatory power. As most existing *C. elegans* mutations are transitions due to EMS mutagenesis bias (Flibotte *et al*. 2010), we first focused on differentiating between guanine (G) to adenine (A) SNSs. While all primers designed to distinguish between G to A transitions had similar melting temperatures, we found that genetic context affected specificity (Figure 1A-D). ARMS primers were able to discriminate between wild type and *him-5(e1490)* across the entire gradient (Figure 1A). However, primers designed to distinguish between wild-type and the *lev-11(x12), cil-7(gk688330)*, and *klp-6(sy511)* alleles were only discriminatory at the highest annealing temperatures when identical concentrations of clean genomic DNA were used (Figure 1B-D). We also tested the capability of ARMS primers to distinguish between other variants such as thymine (T) to A in *plx-2(gk2864)* and G to T in C05B5.11*(gk2895)*. While the primers that detect the *plx-2* and C05B5.11 mutant alleles were specific across the entire temperature range, the wild-type detecting primers exhibited only weak selectivity at high annealing temperatures (Figure 1E,F).

**Figure 1.**
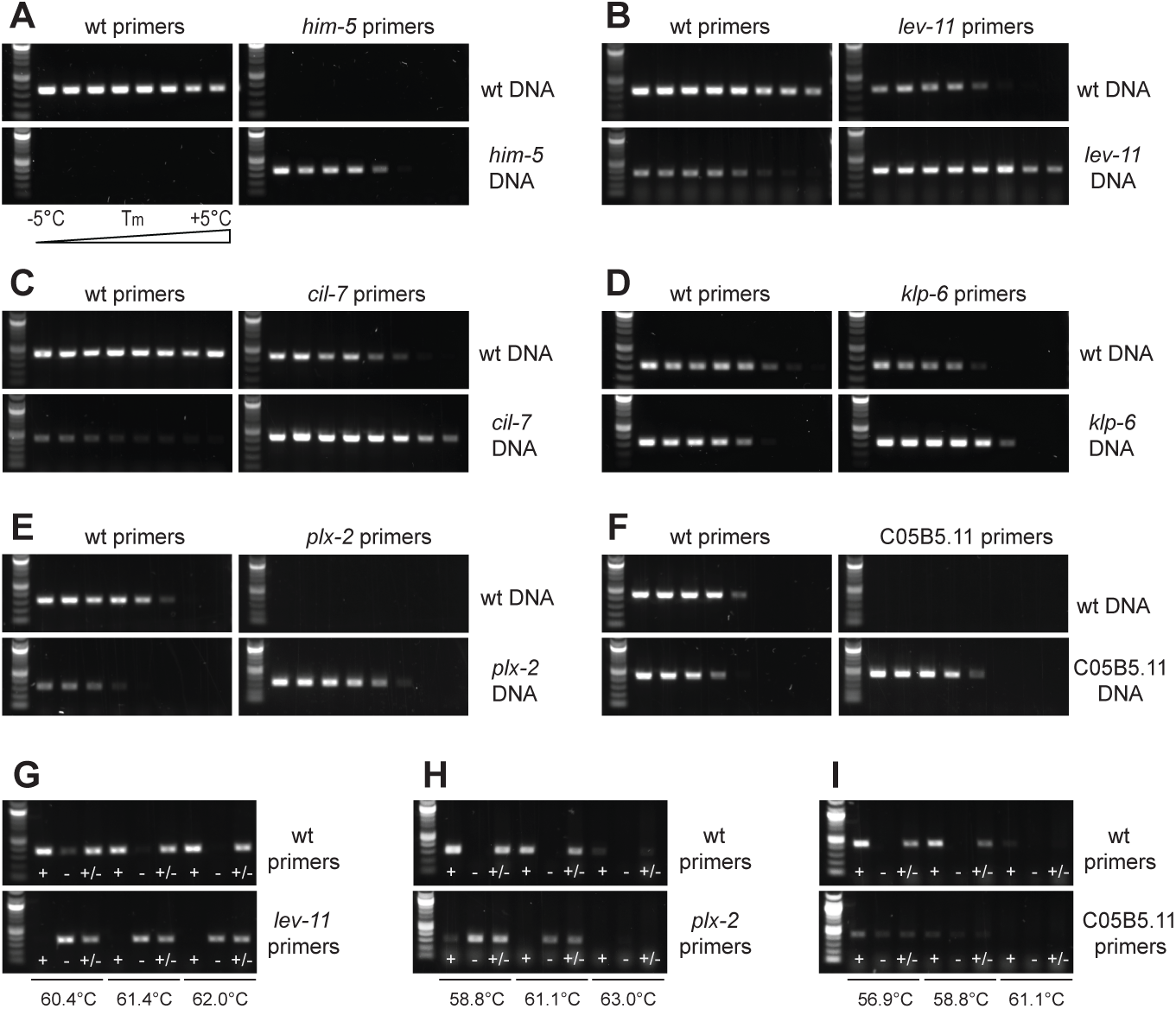
Genotyping mutants with ARMS primers. (A-F) Gradient PCR shows varying specificity of ARMS primers in distinguishing (A) *him-5(e1490)*, (B) *lev-11(x12)*, (C) *cil-7(gk688330)*, (D) *klp-6(sy511)*, (E) *plx-*2*(gk2864)* and (F) C05B5.11*(gk2895)* mutant alleles from the wild type (wt). Gradient temperatures here and throughout were across 10°C; T_m_ minus 5°C (lowest temperature) to T_m_ plus 5°C (highest temperature) as shown in (A). The T_m_ for each primer is indicated in Supplementary Table 1. (G-I) PCR performed on crude *C. elegans* lysate from wild type (+), mutant (-), and heterozygous animals (+/-) at three temperatures optimal for specificity based on gradient PCR. The wild type can be distinguished from (G) *lev-11(x12)* and (H) *plx-2(gk2864)*, but not (I) C05B5.11*(gk2895)*.

We next determined if ARMS primers could be used for routine genotyping with DNA from crude worm lysis, which is of lower quality and contains PCR inhibitors. Using annealing temperatures optimal for specificity based on the gradient PCRs, the wild type could be distinguished from the *lev-11* and *plx-2* mutants as well as heterozygotes over a small temperature interval (Figure 2G,H). However, it was not possible to distinguish the C05B5.11 mutant from the wild type or heterozygote because at temperatures required for specificity, amplification efficiency was low (Figure 1I). These results demonstrate that the ARMS PCR genotyping method requires extensive experimentation to identify the optimal annealing temperature and cannot always be used to distinguish between alleles. Furthermore, there is no flexibility in the placement of ARMS primers, which prevents the use of this method for genotyping alleles in difficult genetic contexts.

**Figure 2.**
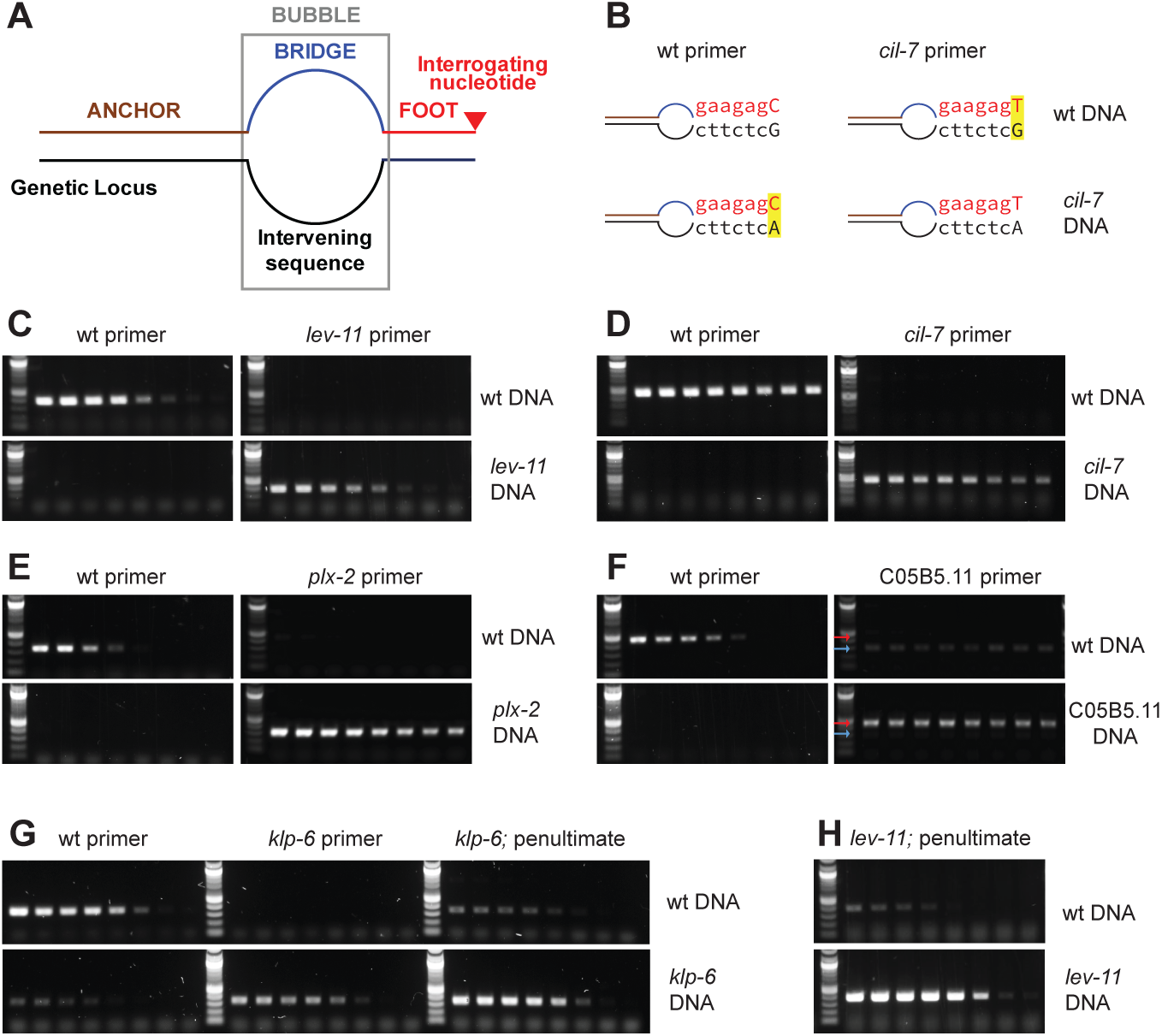
SS primers exhibit specificity across a broad range of annealing temperatures. (A) Each SS primer contains a 5’ anchor sequence (brown), a bridge sequence (blue) not complementary to the template (black), and a 3’ foot sequence (red) that is perfectly complementary for one allele, but contains a mismatch at the interrogating nucleotide (red triangle) for the other allele. The bridge and intervening sequences form a bubble (grey box). (B) Four annealing schematics to illustrate SS primers used to detect wild type and *cil-7(gk688330*) in the presence of wild-type and mutant DNA. In (1) and (4) there is perfect complementarity between the primer and template. In (2) and (3) a mismatch (yellow) at the interrogating nucleotide (capital letter) results in unstable pairing between the primer and template. (C-F) SS primers with an anchor T_m_ of close to 60 °C, 14 bp bridge and 7 bp foot discriminate between wild type and (C) *lev-11(x12)*, (D) *cil-7(gk688330)*, (E) *plx-2(gk2864)*, and (F) C05B5.11*(gk2895)* across the entire gradient PCR. In (F), C05B5.11*(gk2895)* mutant (red arrow) and non-specific (blue arrow) amplification are indicated. (G) The SS primer to detect the wild type allele from the *klp-6(sy511)* mutant allele is not completely selective at low annealing temperatures; the *klp-6(sy511)* mutant SS primer exhibits perfect selectivity. Placement of the interrogating nucleotide at the penultimate position increases PCR efficiency, but decreases specificity of (G) *klp-6(sy511)* and (H) *lev-11(x12)* mutant SS primers.

### SuperSelective primers exhibit discriminatory power for PCR genotyping

We searched for an alternative genotyping method for point mutants and discovered SS primers, which had previously been used for detection of rare variants in qPCR assays (Vargas *et al*. 2016). A SS primer contains a long 5’ sequence termed the “anchor” which anneals to the template and is separated from a short 3’ “foot” sequence complementary to the region around the mismatch by a “bridge” which is not complementary to the template intervening sequence (Figure 2A,B). When the primer is hybridized to the template, the bridge and intervening sequence in the template form a bubble that separates the anchor from the foot. The terminal 3’ nucleotide in the foot, termed the “interrogating nucleotide,” distinguishes the allele variant. Because the foot is short, even one mismatch destabilizes binding and primer extension cannot occur.

To test if SS primers could be used for end point PCR to distinguish *lev-11(x12)* from wild type we designed two allele-discriminating forward primers, one for wild type and the other for the *lev-11* mutant following the rules described by Vagas et al. Each SS primer had an anchor with a melting temperature (T_m_) of approximately 60°C, a 14 base pair (bp) bridge, and a 7 bp foot with the interrogating nucleotide located at the 3’ end. As performed with the ARMS primers, we set up two sets of PCR reactions in parallel for each genomic DNA. One PCR reaction contained the wild-type primer with a common reverse primer, while the other contained the mutant allele-specific primer with the common reverse primer. We observed a dramatic increase in discriminatory ability of SS primers compared to the ARMS primers as the SS primers that detected the wild-type and *lev-11* mutant alleles were perfectly selective across a wide range of gradient temperatures (Figure 2C; compare to Figure 1B). *cil-7, plx-2*, and C05B5.11 mutants, which were poorly distinguished from the wild type with ARMS primers, were also successfully discerned with SS primers across all annealing temperatures (Figure 2D-F). While the SS primer used to distinguish the wild type from the *klp-6* mutant allele did not exhibit complete specificity (Figure 2G), there was significant improvement compared to the ARMS primer (Figure 1D). These results show that SS primers can be used to detect SNSs in different genetic contexts over a broad range of annealing temperatures.

Taq polymerase exhibits less efficient amplification when primers contain an A or T on the 3’ end instead of a G or cytosine (C). Since Vargas et al. found that positioning the interrogating nucleotide at the penultimate position did not affect specificity, we added a C, complementary to the template, to the 3’ end of the *klp-6* and *lev-11* mutant primers, which have an A and T at the interrogating nucleotide, respectively. While this did increase amplification efficiency, it reduced discriminatory power (Figure 2G,H). Thus, to distinguish between wild type and a transition mutant, the mismatch should be placed at the 3’ terminal nucleotide.

### Manipulating the bridge region of the SS primer increases efficiency

Having established the use of SS primers for end point PCR genotyping, we next sought to probe different regions of the primer to develop simple rules for design. All SS primers in Figure 2 contained a 14 bp bridge, with the corresponding intervening sequence also 14 bp, forming a symmetrical bubble. To determine minimum bridge length, we investigated how bubble circumference impacts SS primer efficiency and specificity. We designed additional SS primers to distinguish the wild type from the *klp-6* mutant allele, each with an anchor T_m_ of ∼60°C and 7 bp foot sequence, but different symmetrical bubbles. Comparison of SS primers with 6 bp, 8 bp, and 14 bp bridge sequences for both the wild-type and *klp-6* mutant alleles showed that the smallest bubble circumference resulted in the greatest amplification efficiency (Figure 3A,B). However, the wild-type SS primer containing the 6 bp bridge sequence was non-specific across all annealing temperatures (Figure 3B). SS primers with 6 bp, 8 bp, and 14 bp bridge sequences maintained specificity in distinguishing the wild-type from the *cil-7, plx-2*, and C05B5.11 mutants, with the SS primers containing a 6 bp bridge exhibiting the greatest efficiency (Figure 3C-E). This suggests that irrespective of the 3’ mismatch and genetic context, smaller bubble circumference corresponds with an increase in amplification.

**Figure 3.**
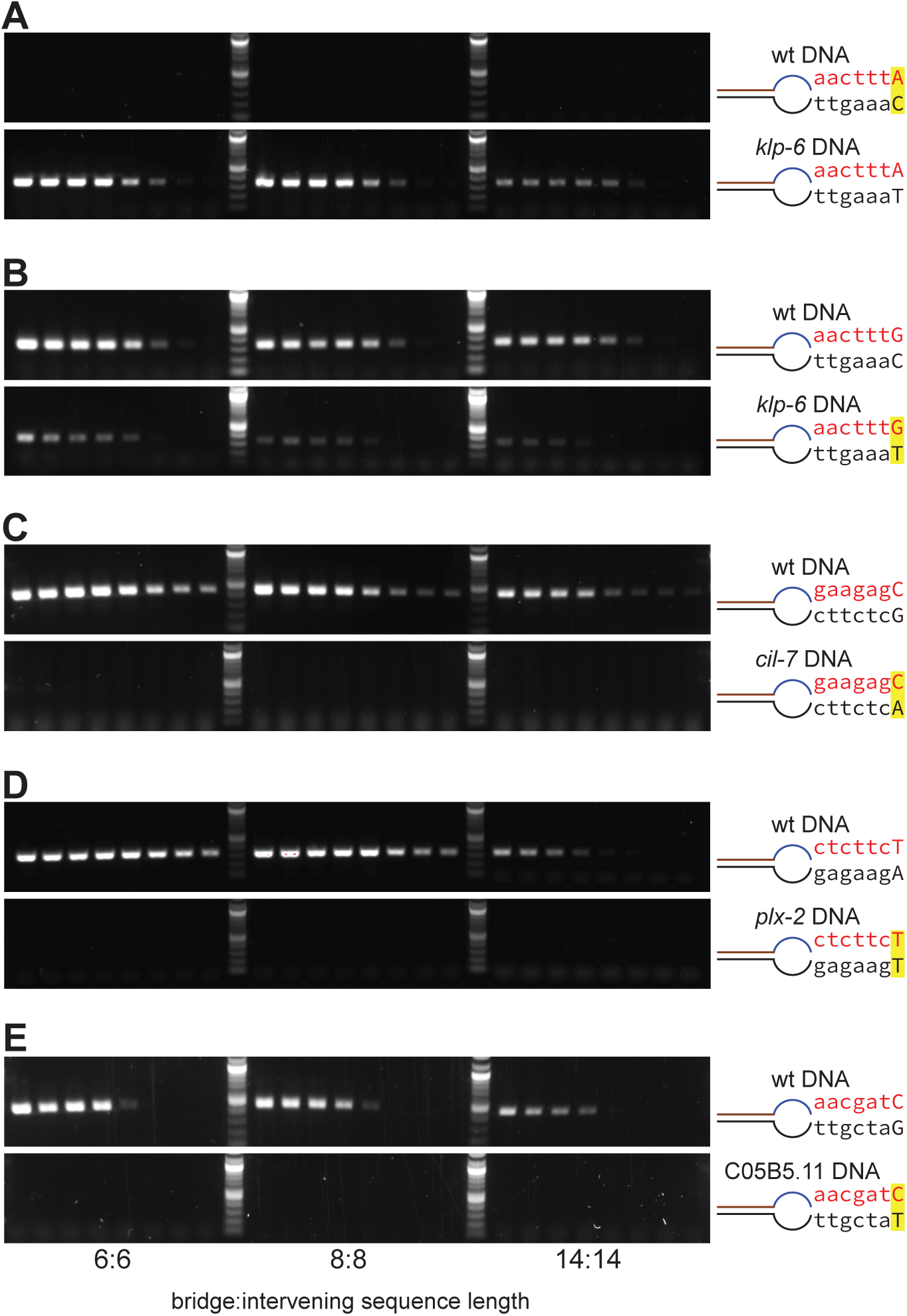
A decrease in bubble circumference increases PCR efficiency. (A-E) Gradient PCR with SS primers to detect the (A) *klp-6(sy511)* mutant allele and wild-type alleles at the (B) *klp-6*, (C) *cil-7*, (D) *plx-2*, and (E) C05B5.11 loci. SS primers have an anchor T_m_ of 60°C, 7 bp foot and varying bridge sequence length (6 bp, 8 bp, and 14 bp); length of intervening sequence is equal to the length of bridge sequence, creating a symmetrical bubble. Primer schematics as in Fig. 2; yellow highlight indicates mismatch.

The wild-type SS primer, which forms a G-T mismatch with the *klp-6* mutant sequence was less specific than all other SS primers tested (Figure 3B). This purine-pyrimidine mismatch has a similar geometry to G-C and A-T base pairings, causing only a weak destabilizing effect, which enables it to be extended more efficiently by Taq polymerase than any other mismatch (Huang *et al*. 1992; Rejali *et al*. 2018). To determine whether the non-specificity of the SS primer used to detect the wild-type allele at the *klp-6* locus was due to genetic context or the weak G-T primer-template mismatch, we designed SS primers with short bridge sequences to distinguish the *lev-11(x12)*, daf-7*(e1372)*, and *eat-2(ad465)* G to A transition mutants from wild type. SS primers with a 6 bp bridge sequence corresponding to a 6 bp template intervening sequence specifically detected the wild type allele across all annealing temperatures at the *daf-7*, but not *lev-11* and *eat-2* mutant loci. However, specificity was lost when the bridge sequence was shortened to 4 bp for all G-T primer-template mismatches (Figure 4A-C). SS primers used to discriminate the wild-type allele from *plx-2(gk2864)*, C05B5.11*(gk2895)* and *lev-11(x12)*, which result in T-T, C-T and C-A primer-mutant template mismatches respectively, were specific even with a short 4 bp bridge (Figure 4D-F). These results show that the minimum circumference of the bubble needed to maintain specificity depends on both primer-template mismatch and genetic context.

**Figure 4.**
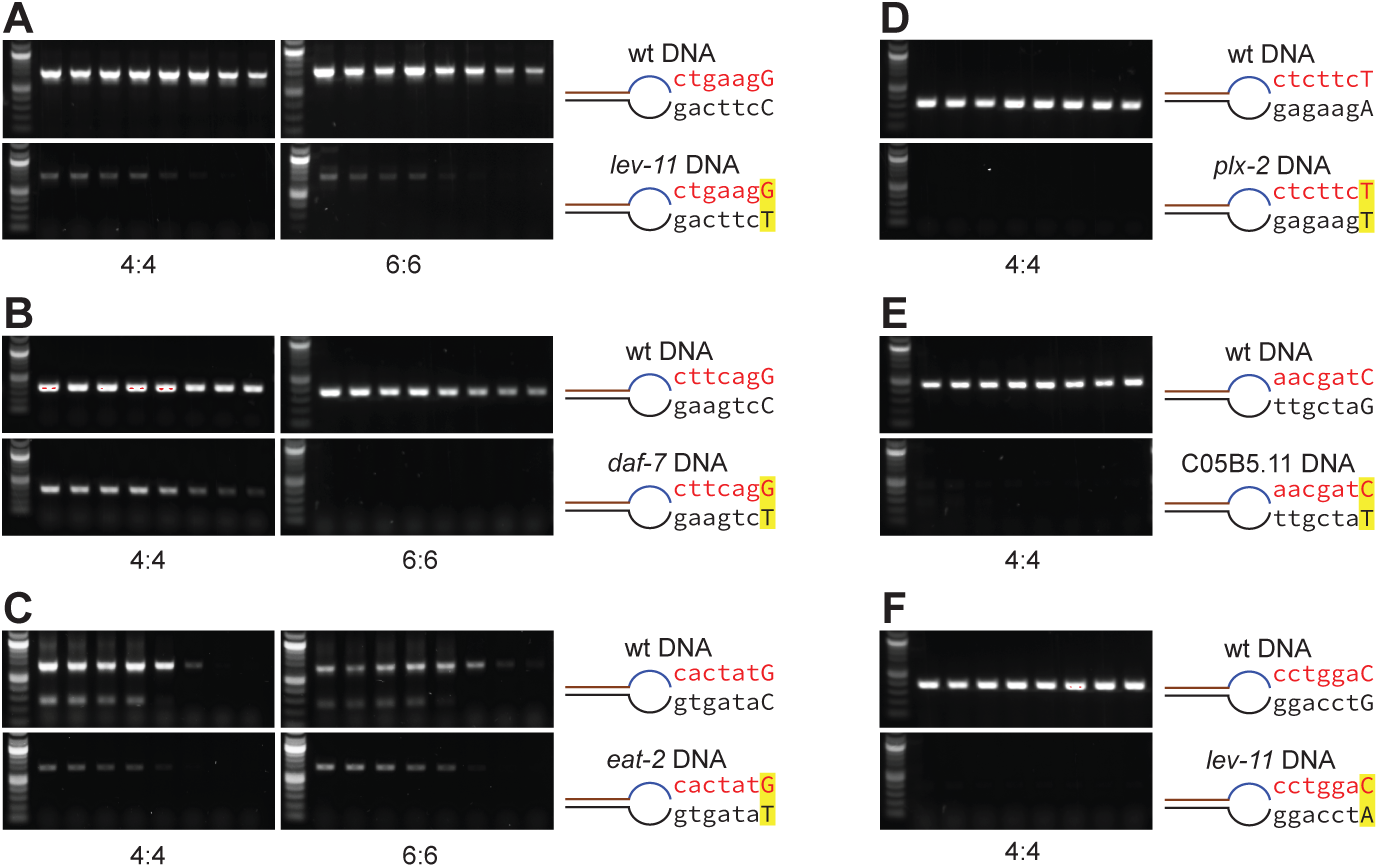
Guanine-thymine primer-template mismatches are the least discriminatory. (A-C) SS primers with an anchor T_m_ of 60 °C, 7 bp foot, and either 4 bp or 6 bp bridge sequence cannot perfectly distinguish the wild type from (A) *lev-11(x12)*, (B) *daf-7(e1372)*, and (C) *eat-2(ad465)* mutant alleles. The SS primer-template mismatch at the interrogating nucleotide is G-T (yellow); intervening sequence length is equal to bridge sequence length (4:4 and 6:6). (D-F) SS primers with a 4 bp bridge sequence exhibit complete specificity when interrogating nucleotide mismatch is (D) T-T in the *plx-2(gk2864)* mutant, (E) C-T in the C05B5.11*(gk2895)* mutant, and (F) C-A in the *lev-11(x12)* mutant. The SS primer to detect the wild-type from *lev-11(x12)* mutant allele in (F) anneals to the opposite strand compared to (A).

### The foot region of the SS primer can be manipulated to increase specificity

We next investigated how the length of the foot region impacts efficiency and specificity using SS primers that detect the wild-type allele at the *klp-6* locus. Our original 14:14 SS primer contained a 7 bp foot sequence with the interrogating nucleotide on the 3’ end (Figure 2G). We discovered that shortening the foot sequence to 5 or 6 bp decreased efficiency without affecting specificity (Figure 5A). A SS primer with a 4 bp foot sequence did not produce any product (Figure 5A) even at a low 45° C annealing temperature (data not shown).

**Figure 5.**
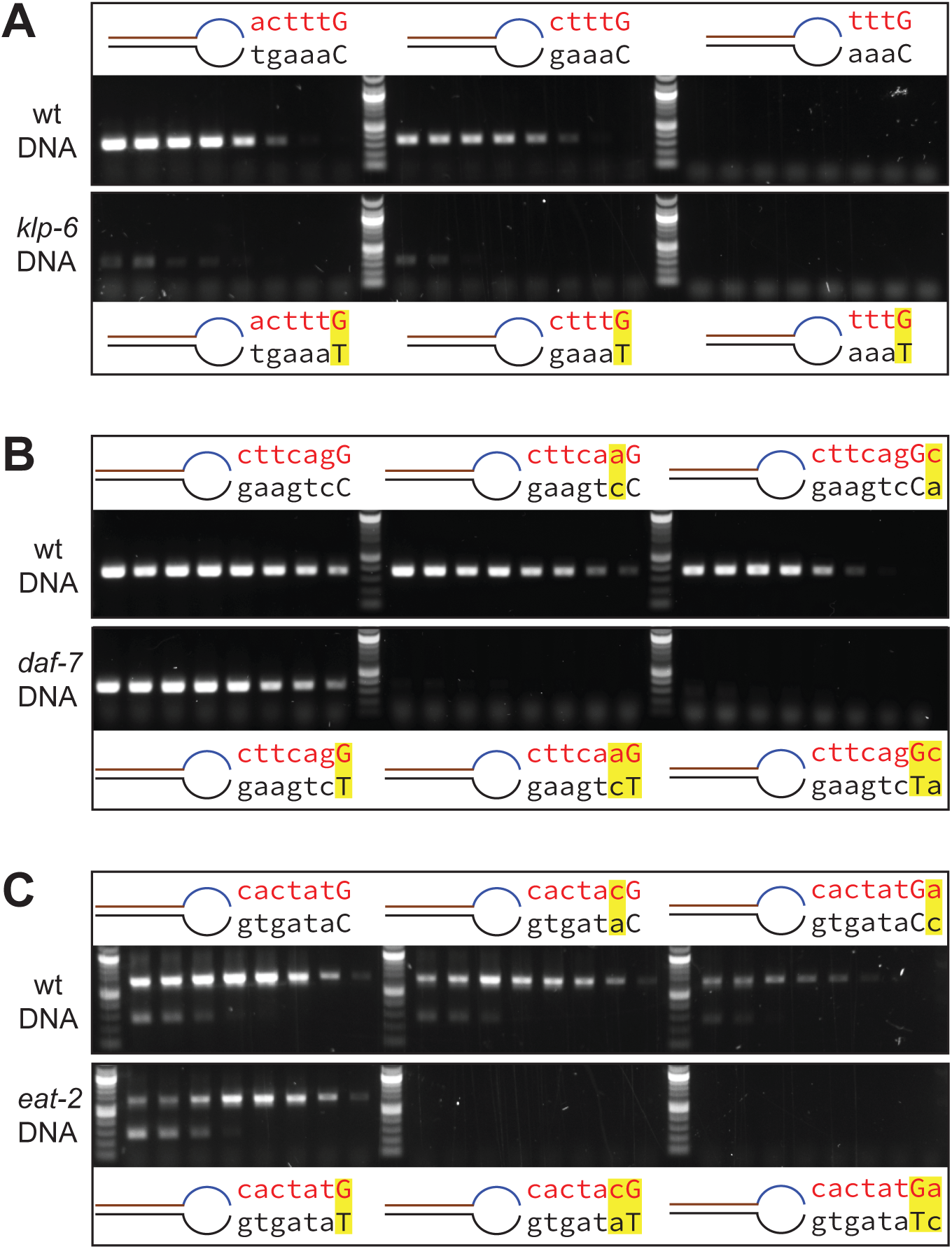
SS primer foot sequence and length influence specificity and efficiency. (A) Decreasing foot length from 6 bp (left) to 5 bp (middle) decreases efficiency; a 4 bp foot (right) eliminates amplification. Primers detect the wild-type allele at the *klp-6* locus; bridge is 6 bp. (B-C) Introduction of a weak purine-pyrimidine mismatch at the penultimate (middle) or the terminal position (right) increases specificity of SS primers with a 4 bp bridge that detect the wild-type allele at the (B) *daf-7* and (C) *eat-2* loci. In primer schematics, the interrogating nucleotide is capital, mismatches highlighted in yellow.

Since shortening the foot sequence had an undesirable effect on amplification, we sought to determine if additional mismatches in the foot sequence could be used instead to increase SS genotyping specificity. We introduced a mismatch at the penultimate position to the interrogating nucleotide, which we designate the (−1) position. Placing a G-A mismatch, which has a strong destabilizing effect (Rejali *et al*. 2018), at the (−1) site prevented amplification (Supplementary Figure 1). However, introduction of a weak A-C purine-pyrimidine mismatch at the (−1) position in SS primers with 4 bp bridge sequences that previously could not distinguish wild type from *daf-7* and *eat-2* mutant alleles, resulted in specificity across all annealing temperatures (Figure 5B,C). Likewise, introduction of a purine-pyrimidine mismatch terminal to the interrogating nucleotide also generated specificity (Figure 5B,C). This suggests that placement of an additional weak destabilizing mismatch in the foot can be used to increase discriminatory power.

### SS primers enable flexibility in anchor placement

In primer design, it is important to avoid runs of one base, A/T rich domains, tandem repeats, and sequences that form secondary structure. We considered that changing the length of the template intervening sequence would allow for anchor placement flexibility. To determine how amplification is affected by an asymmetric bubble, we designed a SS primer with a 6 bp bridge to a 24 bp intervening sequence (6:24) and a 7 bp foot to distinguish wild type from the *lev-11* mutant and observed specific amplification across the entire gradient (Figure 6A). However, we saw little amplification when SS primers with 6:24 and 6:30 asymmetric bubbles were used for detection of the wild-type allele at the *klp-6* and *cil-7* loci, respectively (Figure 6B,C). Increasing foot length to 8 bp, with a C in the 3’ terminal position improved efficiency of SS primers with asymmetric bubbles without affecting discriminatory power (Figure 6B,C). Since the sequence surrounding *daf-7(e1372)* is A/T rich, we created a SS primer that forms an extremely asymmetric 6:51 bubble, and this primer perfectly discriminated the wild-type from the *daf-7* mutant allele across all gradient temperatures. These results demonstrate that the anchor of SS primers can be moved to enable genotyping in difficult genetic contexts.

**Figure 6.**
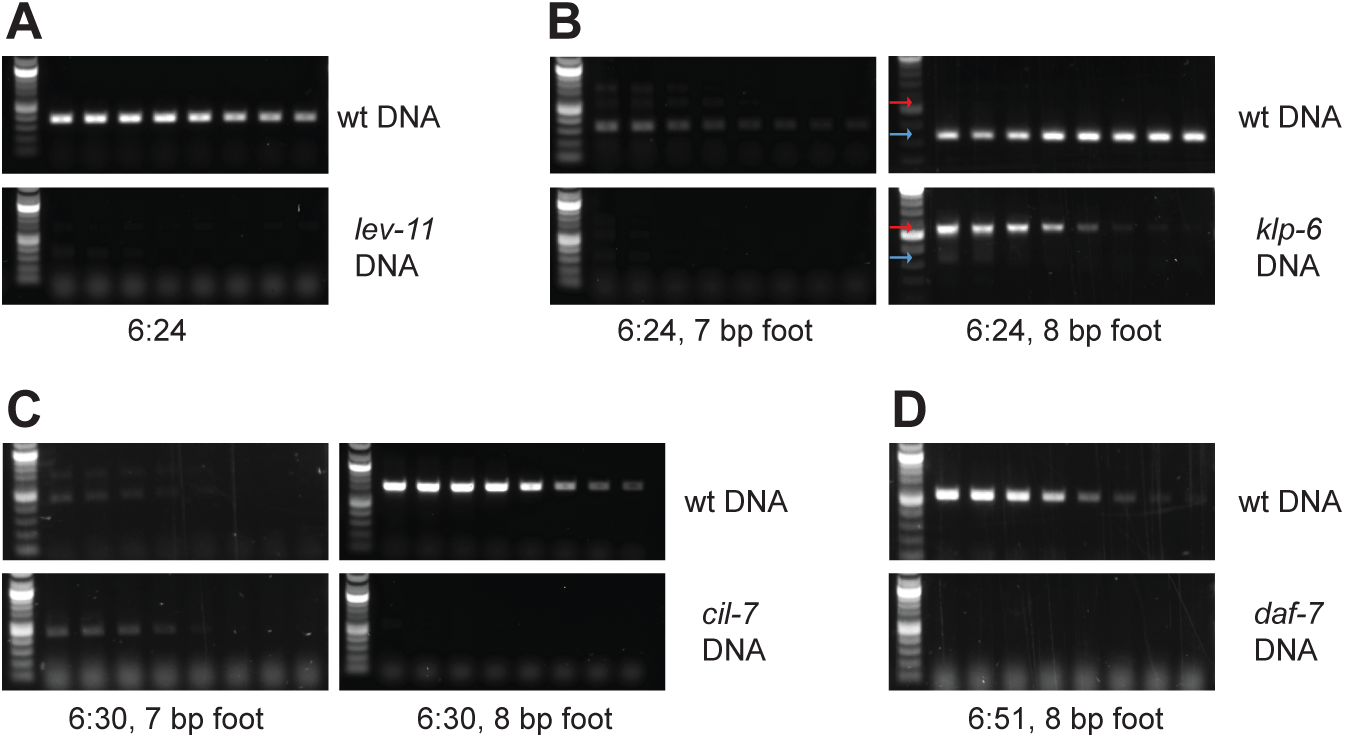
Asymmetric bubble design allows for flexibility in anchor placement. (A) A SS primer with a 6 bp bridge, which forms an asymmetric bubble with a 24 bp template intervening sequence (6:24), detects the wild-type allele at the *lev-11* locus. (B,C) Asymmetric bubbles result in poor amplification (left panels); efficiency is improved by increasing the length of the foot. In (B) wild-type (blue arrow) and non-specific (red arrow) amplification are indicated. (D) A SS primer which forms a 6:51 asymmetric bubble with the template detects the wild-type allele at the *daf-7* locus.

## DISCUSSION

We developed a rapid, low-cost method for detection of point mutants by optimizing SS primers for end point PCR. Our analyses of seven separate genetic contexts and eight different types of mismatches show that SS primers can be used universally for genotyping over a broad range of annealing temperatures. We discovered that balancing stabilizing versus destabilizing factors in the foot region affects specificity, while decreasing bridge length increases efficiency. Amplification occurs even when the SS primer bridge and intervening template sequence form an asymmetric bubble, allowing for flexibility in anchor placement. Our work demonstrates the power of SS primers for routine genotyping and we propose that this method could also be used for SNP mapping, screening of CRISPR mutants, and identification of site-directed mutagenesis clones through colony PCR.

### Simple instructions for SS primer design

We have defined several important factors to consider when carrying out SS genotyping. To design a SS primer, first identify a 7 base pair foot with the interrogating nucleotide in the terminal 3’ position. Second, identify a 5’ anchor sequence with a 50 to 60% G/C content and a T_m_ ∼60 °C at least 6 bp away from the foot. In many cases, a symmetric bubble consisting of a 6 bp bridge between the anchor and foot in the primer and corresponding non-complementary 6 bp intervening sequence in the template provides both good efficiency and specificity. However, the intervening sequence length can be increased to enable placement of the anchor in a more favorable position. Third, consider the mismatch between the primer and template at the interrogating nucleotide. A weak G/T mismatch will reduce the ability to detect between the target and non-target allele. To decrease undesired stability between the primer and non-target template, a second mismatch can be introduced either penultimate or terminal to the interrogating nucleotide. Fourth, design another SS primer to detect the other allele as well as a common reverse primer. Finally, make sure that the SS primers do not have secondary structure using the IDT OligoAnalyzer. While we have used gradient PCR to examine the properties of SS primers, given that specificity is generally observed across the entire gradient, we recommend an annealing temperature of 58°C for routine genotyping. No more than 30 cycles should be used since the number of amplicons produced by the perfectly matched primer should reach plateau by this point, and if the PCR runs for additional cycles, undesired products will continue to be amplified exponentially (Saiki *et al*. 1988).

### Effect of specific primer-template mismatches on PCR specificity and efficiency

A single 3’ terminal mismatch destabilizes primer-template interaction, and as *Taq* DNA polymerase does not possess 3’ to 5’ exonuclease activity for mismatch repair, this mismatch reduces extension efficiency, and as a result, PCR amplification, when compared with a primer perfectly complementary to the template (Petruska *et al*. 1988; Tindall and Kunkel 1988; Huang *et al*. 1992; Rejali *et al*. 2018). While this serves as the foundation for allele-specific detection with SS genotyping, PCR amplification is also influenced by the specific primer-template mismatch, with purine-purine mismatches being the most inhibitory, and purine-pyrimidine mismatches being the least inhibitory (Huang *et al*. 1992; Rejali *et al*. 2018). Ethyl methanesulfonate (EMS), the primary chemical mutagen used for forward genetic screens in *C. elegans*, exhibits a mutagenesis bias toward G/C to A/T transitions (Flibotte *et al*. 2010). When differentiating between EMS generated alleles, a G at the interrogating nucleotide of the wild-type detecting primer mismatches with a T in the mutant template. Here we found that primers with a G-T mismatch were less selective than those with T-T, C-T and C-A mismatches, consistent with the G-T mismatch being the least inhibitory (Huang *et al*. 1992; Rejali *et al*. 2018). As previously reported for extension rate (Huang *et al*. 1992), we observed that sequence context influenced end point PCR genotyping for weak G-T mismatches.

To decrease extension efficiency, and thus improve PCR specificity, an additional mismatch can be introduced either penultimate or terminal to the interrogating nucleotide (Ugozzoli and Wallace 1991; Bui and Liu 2009). Some purine-purine penultimate mismatches such as G-A inhibit extension efficiency even more than a 3’ terminal G-T mismatch (Rejali *et al*. 2018). In fact, we found that a G-A mismatch at the penultimate position in the SS primer to detect the wild-type allele at the *klp-6* locus prevented amplification. Thus, if introducing an additional mismatch at the penultimate position, strong G-A, G-G, A-A, and C-C primer-template mismatches should be avoided, while weak G-T and C-A mismatches are tolerated.

### SS genotyping offers distinct advantages compared to other methods

Here we consider how SS primers compare with other existing allele discrimination methods. Mutations that result in creation or disruption of a restriction site can be detected by amplification of the template from the target region followed by enzymatic digestion of the DNA and electrophoresis. However, genotyping of many different alleles by this method requires a large collection of different restriction enzymes and a suitable restriction enzyme or artificial restriction site cannot be introduced at all locations. SS genotyping can be used to distinguish between mismatches in all genetic contexts and does not require any reagents or effort beyond PCR. Further, unlike single-base extension genotyping (Sauer 2000; Trewick *et al*. 2011), the 5’ fluorogenic nuclease Taqman assay (Livak *et al*. 1995; Callegaro *et al*. 2006), and melting curve analysis of FRET probes (Livak 1999; Combrinck *et al*. 2013), expensive equipment and specialized training are not required for design and use of SS primers for allele detection.

Similar to SS genotyping, ARMS PCR and the modified simple allele-discriminating PCR are inexpensive methods which utilize allele-specific oligonucleotide primers (Little 2001; Bui and Liu 2009; Medrano and De Oliveira 2014). However, when genotyping *C. elegans* point mutants, we found that ARMS PCR required extensive troubleshooting to determine optimal annealing temperature and could not always be used to distinguish between alleles at any temperature. When genotyping with crude *C. elegans* DNA lysates, PCR amplification can be affected by variability in lysis efficiency and DNA concentration if different numbers of worms are used. Given that stringent reaction conditions are required for allele discrimination with ARMS PCR, we remain concerned that the quality of the starting template and small fluctuations in temperature could impact genotyping results. Further, unlike SS primers, there is no flexibility in placement of ARMS and SAP primers, which makes allele detection difficult in certain genetic contexts (Medrano and De Oliveira 2014). In conclusion, SS genotyping is 1) low cost, 2) does not require special equipment, 3) works over a broad range of annealing temperatures, and 4) allows for flexibility in primer placement. SS primers can theoretically be utilized in all organisms and for any laboratory applications that require discernment between alleles.

## ACKNOWLEDGEMENTS

We thank the *Caenorhabditis* Genetics Center (CGC) for strains and Aimee Jaramillo-Lambert and members of the Tanis lab for critical reading of the manuscript. This work was supported by a National Institutes of General Medical Sciences (NIGMS) IDeA Network of Biomedical Research Excellence (INBRE) P20 GM103446 Pilot Project grant (to J.E.T).

## FIGURE LEGENDS

**Supplemental Figure 1.**
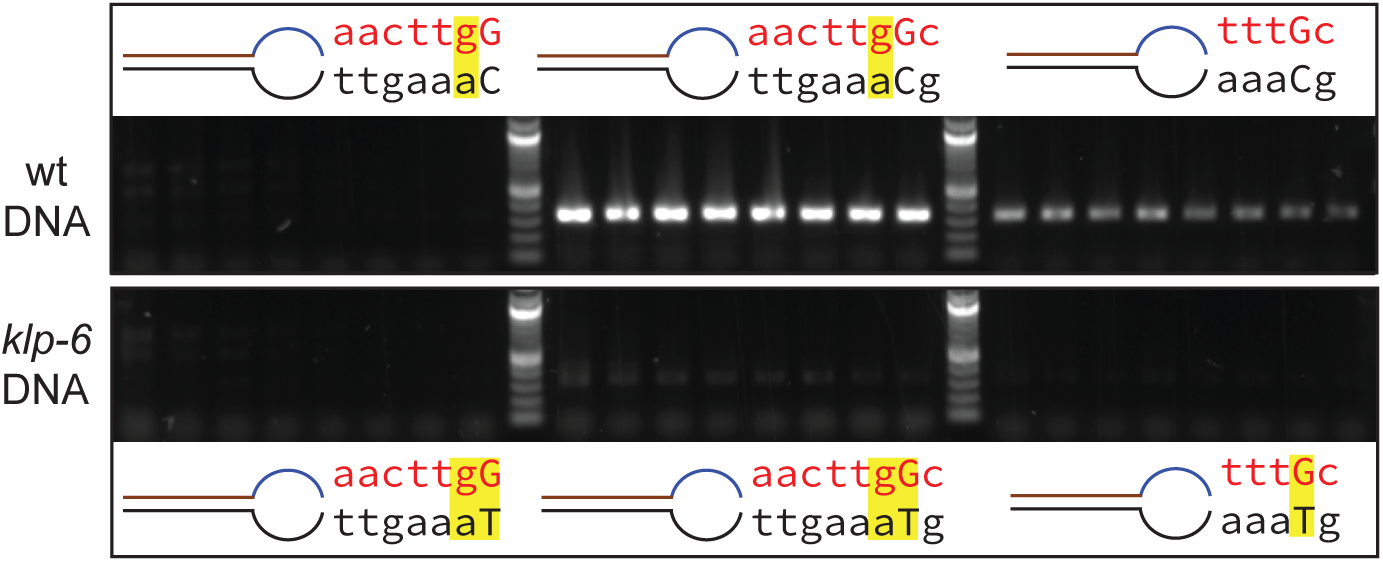
Placement of the interrogating nucleotide at the penultimate position increases efficiency. Introduction of a strong purine::purine mismatch (−1) to the interrogating nucleotide prevents amplification (left), which can be restored by adding an additional complementary terminal base (middle). Placement of the interrogating nucleotide in the penultimate position in a SS primer with a short foot increases efficiency (compare to middle and right hand panels of Fig. 5A). All primers here detect the wild-type allele at the *klp-6* locus. In the schematics, the interrogating nucleotide is in capital, all mismatches highlighted in yellow.

**Supplementary Table 1:**
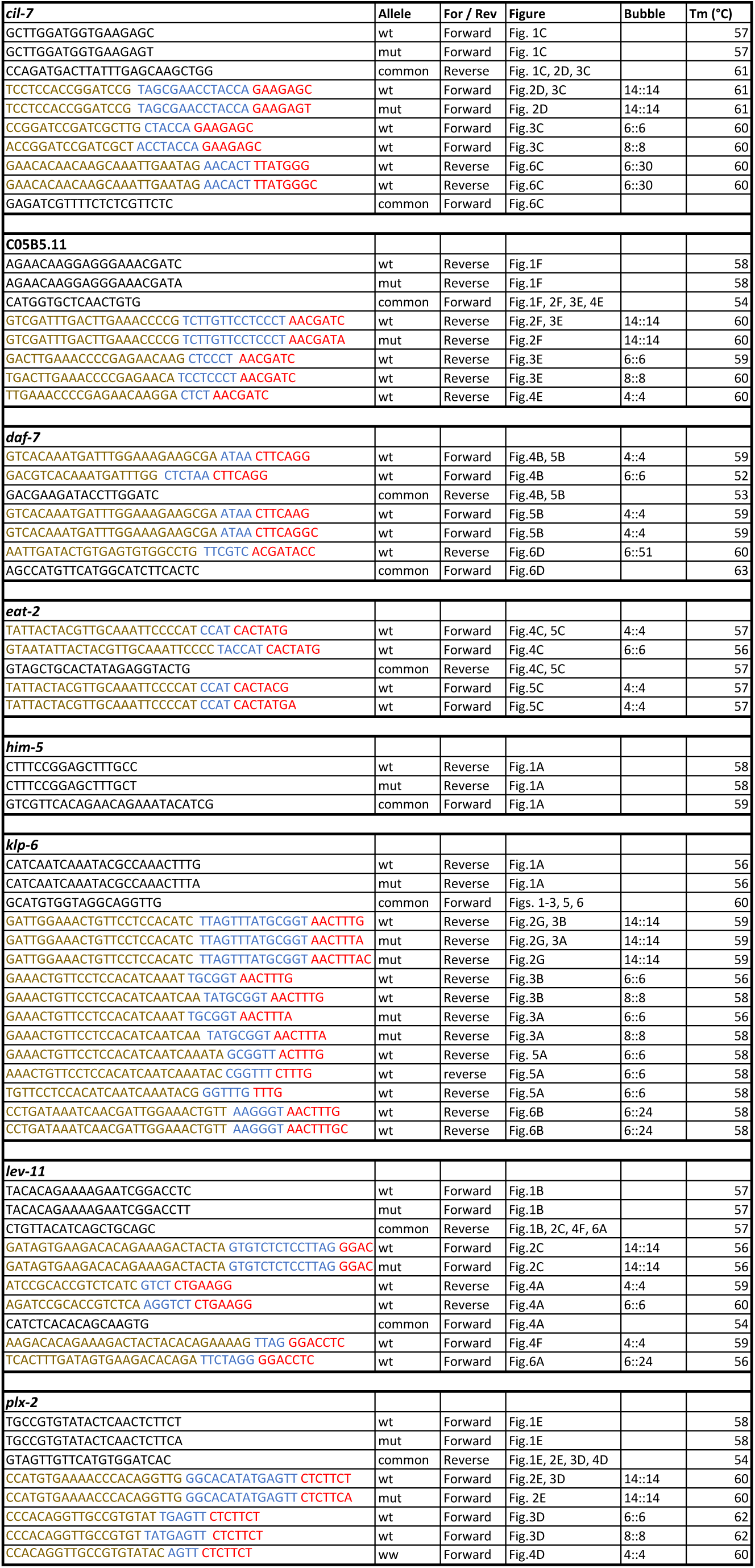
For SS primers, the 5’ anchor sequence (brown), bridge sequence (blue), and 3’ foot sequence (red) are indicated

